# Continuous Monitoring of Head Turns: Compliance, Kinematics, and Reliability of Wearable Sensing

**DOI:** 10.1101/2025.04.29.650670

**Authors:** Selena Y. Cho, Leland E. Dibble, Peter C. Fino

## Abstract

Wearable devices offer objective mobility metrics for continuous monitoring but often focus on traditional measures like step count or gait speed. Other quantitative metrics such as head kinematics may provide valuable insights into mobility, balance, and sensory integration, given the head’s central role in coordinating vestibular, ocular, and postural control. Yet, basic knowledge about capturing daily living head turns, including participant compliance, algorithms, normative data, and reliability, is not yet established. This study aimed to resolve this knowledge gap by capturing head and trunk movement kinematics over a 7-day period and to establish normative data in healthy adults. Participants (n = 24) wore head-mounted sensors for an average of 16.38 hours per day (SD = 4.43), completing 5,163 (SD = 1,466) head turns daily, with 72% occurring independently of trunk motion. Head turn amplitude (M = 58.18°, SD = 4.26°) was comparable to lumbar turns, while peak velocity was higher for head turns (M = 104.49°/s, SD = 12.08°/s). By the second day, all head turn metrics achieved excellent reliability (ICC > 0.9), supporting the feasibility of multi-day monitoring. Additionally, we examined the relationship between head motion and other mobility metrics and established recommendations for implementing similar protocols for capturing future studies, including the minimum number of days required for reliable data collection. Findings from this study provide a foundation for future multi-day continuous monitoring of head kinematics in both healthy and clinical populations.

## I. Introduction

OBJECTIVE measures of mobility from wearable devices are increasingly used for continuous monitoring of patients during free-living daily life [1-4]. Common measures such as step count, gait speed, and activity level provide insight into the quantity or intensity of physical activity [5]. While these quantitative measures offer macroscopic views of daily function, focusing on the kinematics of individual movements can provide micro-level measures that highlight the quality of movement. In clinical populations, these micro-level measures, such as stride length, turning speed, joint range of motion, and postural stability, often reveal important differences undetected by macro-level outcomes [6-13]. For instance, step count is commonly used to quantify the volume of physical activity and its relationship to morbidity and mortality, but it provides no information on movement quality [14]. Information about movement quality can reveal important clinical information in impaired populations; people with Parkinson’s disease have a similar number of steps per day compared to their neurotypical counterparts but exhibit significant differences in micro-level metrics of quality, like steps per turn and stride variability [15-17]. To extract these micro-level metrics of quality, current applications of continuous monitoring focus on lower-limb kinematics, lumbo-pelvic kinematics, and spatiotemporal gait analysis [18-26]. However, movements of other body segments could offer complementary insights into overall mobility and function.

Head kinematics may provide valuable insights into various aspects of a person’s function during daily living, including visual processing and postural adjustments [27-40]. The head serves as the central hub for crucial sensory systems, including the vestibular and ocular systems, which play fundamental roles in maintaining spatial orientation, balance, and gaze stability. As a result, clinical assessments of balance and mobility often include tasks that involve volitional head movements, such as turning one’s head side to side while walking, to examine how individuals decouple their head from the body and the subsequent consequences on mobility [41-47]. Though clinical assessments largely focus on the effects of head movements on gait stability, the kinematics of these volitional head movements can also offer clinically relevant information. For example, people with complete unilateral vestibular loss following vestibular schwannoma resection surgery exhibit slower and smaller volitional head movements compared to healthy controls [48-54].

Previous research provided clinical motivation for capturing head motion in lab- or clinic-based environments, but few studies have examined head kinematics during free-living daily life [32, 51]. Recent advances in wearable inertial measurement units (IMUs), including smaller sensors, longer battery life, and larger onboard storage capacity, have made continuous monitoring of head motion feasible. For example, Hausamann et al. demonstrated the feasibility of capturing head kinematics over a 24-hour period and reported parameters that measure head stabilization (e.g., harmonic ratios, coherence, and phase differences) [32]. Furthermore, Paul et al. examined relevant discrete head movements (i.e., head turns) and the independence of head turns relative to trunk motion during a 15-minute simulated community ambulatory circuit that required participants to navigate through crowded hallways, ascend and descend stairs, and scan for pedestrian and/or vehicular traffic [51]. Despite the ecological relevance of this community ambulation task, simulated tasks may not represent true daily living or variability over the course of a day or across multiple days. Real-world daily variability often requires multiple days (e.g., 2-3 days) of continuous recordings for most mobility metrics (e.g., gait speed, sway) [55]. The key next step is to conduct multi-day recordings of head movements in real-world settings, offering a more comprehensive understanding of head movement patterns in daily life.

Extending prior work, the purpose of this study was to assess the feasibility of capturing the kinematics of head movements in free-living daily life over a 7-day period and to provide normative, healthy adult data on head kinematics, defined as the number, frequency, amplitude, and velocity of head turns, during daily life. Additionally, this study sought to guide future multi-day, continuous monitoring of head kinematics by addressing the following critical points: 1) Assess participant compliance in wearing head sensors during daily life; 2) Identify methods for achieving repeatable sensor orientation across multiple donning and doffing cycles; 3) Establish normative descriptive statistics for head-turning kinematics in healthy adults; 4) Examine the relationship between head motion and other continuous monitoring metrics; 5) Determine the recommended amount of wear time needed to obtain reliable measures of head-turning kinematics.

## II. Materials AND Methods

### A. Participants

Twenty-four healthy adults (15 females / 9 males, mean age = 28.3 years, SD = 6.1 years) participated in the study. All participants provided informed written consent approved by the University of Utah Institutional Review Board (IRB). Participants were included if they were aged between 20 and 50 years, had no history of neurological or musculoskeletal disorders, and were able to ambulate independently. Individuals were excluded if they had any condition that could affect their gait or balance, were pregnant, or had recent injuries requiring medical intervention.

### B. Instrumentation

Three IMUs (Axivity Ax6, Newcastle, UK) were attached to the participant: one behind the right ear just superior to the mastoid process, one on the thoracic spine at the level of the T2 vertebra spinous process, and one on the lumbar region at the level of the L2-L4 vertebra spinous processes. The thoracic and lumbar sensors served as references for head-on-body and whole-body turns, respectively, similar to past work [17, 56, 57]. The sensors were adhered using double-sided 3M 1522 tape (True Tape, Canon City, Colorado) with the thoracic and the lumbar sensor protected with a clear 4×4in Tegaderm bandage (3M, Maplewood, Minnesota). Each IMU was initialized using the open OMGUI software, (Newcastle, UK) sampling at 100 Hz with a tri-axial accelerometer range of +/-8g, and tri-axial gyroscope range of +/-2000 dps.

### C. Procedure

Participants were instructed to wear the sensors continuously for seven days, removing the head sensor before showering and sleeping. The participants were sent home with additional double-sided tape and bandages as well as a tracking log to record time points of when they took the sensors off, placed the sensors back on, and any gross measure of activity (e.g., what activities they completed each day out of a predefined list). Upon replacing the head sensor (i.e., after sleeping or showering), participants were asked to perform a series of calibration movements that included shaking their head side to side three times, nodding their head up and down three times, and jumping three times. These movements were used to identify reattachment timepoints to aid in processing the IMU data and align the sensor axes with the anatomical axes of the head and thoracic.

### D. Data Preprocessing

All data analysis and signal processing were performed using a custom-written MATLAB script (MathWorks, Natick, MA). The sensors occasionally produced repeated samples, which were corrected using an OMGUI function that resampled the signal at 100 Hz and removed duplicated data points. To address temporal drift, which can range from one to three seconds per day, three large vertical taps on the table were performed simultaneously with all three sensors both before and after the data collection period to facilitate synchronization. These vertical taps were identified, and sensors were synchronized using cross-correlation of this motion to remove any additional sensor lag. Following pre-processing synchronization, data were segmented into 24-hour periods from 00:00:00 to 23:59:59.

### E. Data Postprocessing

To reorient the sensor to the global axis whenever the sensor was placed back on the head, we identified the calibration movements (shaking, nodding, jumping) by detecting three sequential peaks exceeding 40°/s in both the vertical yaw (left-right head turns) and anterior-posterior pitch (up-down head turns) directions, followed by three acceleration peaks greater than 10 m/s^2^ (jumping). To qualify as a calibration movement, each successive peak had to occur within 2 seconds from the previous peak. When a calibration movement was successfully detected, the timepoint marking the start of the first head turn was recorded. Accelerometer data during these periods were used to generate a rotation matrix that aligned the vertical axis with Earth’s gravity, pointing downwards. The same rotation vector was applied to the gyroscope data in all three sensors, ensuring consistent alignment [58, 59].

Despite instructions, not all participants completed the calibration procedure for all reapplications of the head sensor. In cases where the calibration movement was not detected, we used an alternative procedure based on large walking bouts consisting of at least 100 steps to align the sensor axes. This method assumed that the participant was upright, and their head was, on average, in a neutral, forward-looking position while walking [58]. The script generated an array by merging calibration movement timepoints with walking bout timepoints to identify periods for sensor reorientation. A rotation matrix was then applied to reorient the sensor to Earth’s gravity [58, 59].

### F. Turn Detection

To detect head turns with respect to the global reference frame (head-in-space turns), we modified a turn detection algorithm previously used to detect whole-body turns from a lumbar-mounted IMU [60]. The angular velocity signal in the vertical yaw direction was filtered using a finite impulse response (FIR) filter, with the impulse response shaped by an Epanechnikov kernel— a symmetric, weighted moving average with an impulse duration of 1.476s [61]. This filter was selected for its ability to enhance signal clarity through localized weighted averaging, as opposed to frequency-specific elimination techniques (e.g., Butterworth filter). This method maintains the fundamental characteristics of the signal’s core structure, allowing for precise identification of subtle variations in rotational speed indicative of head movements. After filtering, the signal was rectified to ensure that all turns, regardless of direction, were combined into a single metric (Figure 1).

**Fig. 1.**
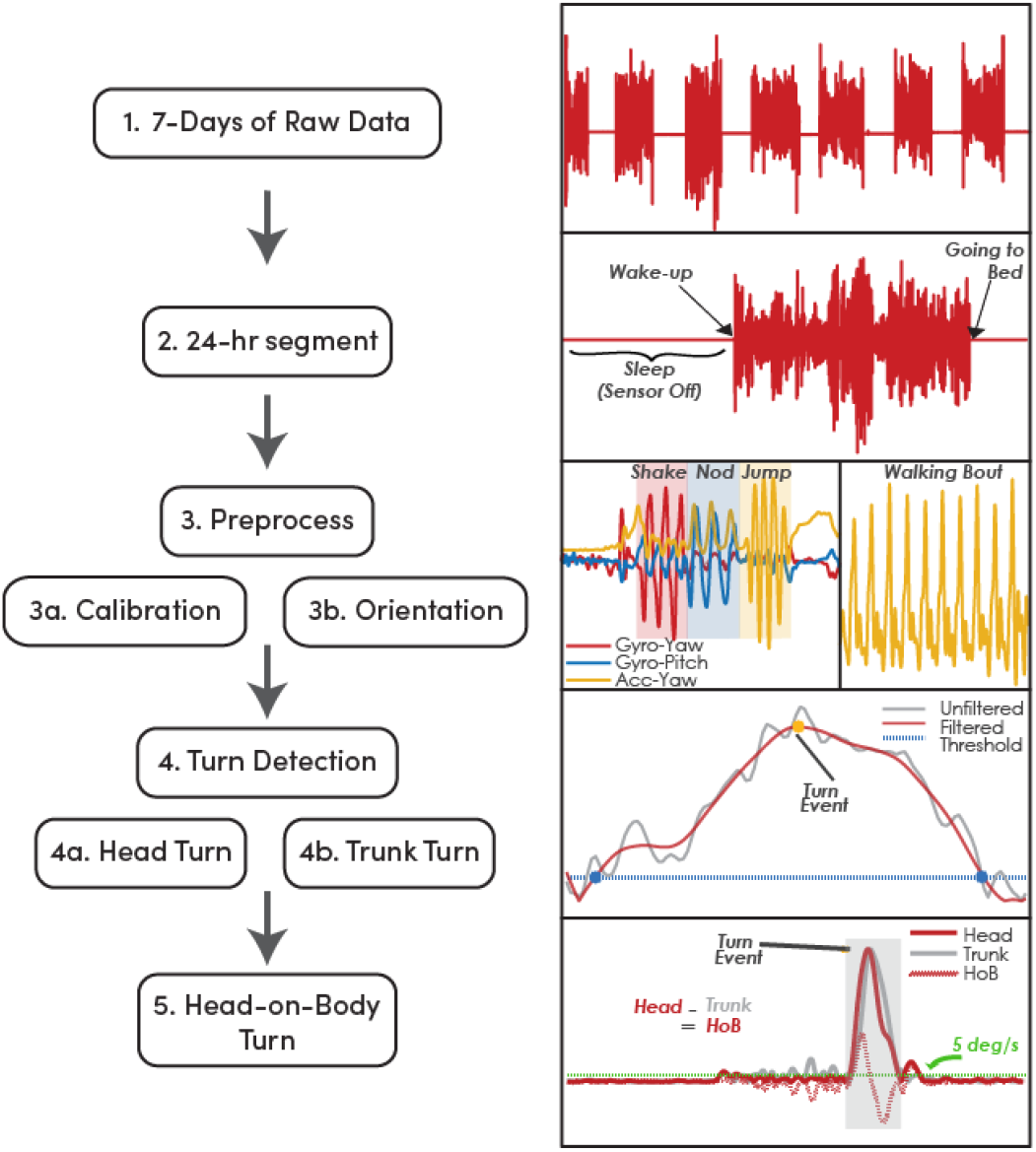
Overview of the data processing pipeline for extracting head and head-on-body turns from wearable sensor data. (1) Seven days of raw data are collected using IMU sensors. (2) Data are segmented into 24-hour periods (3) Preprocessing involves filtering and sensor alignment, including (3a) identifying calibration movements such as head shaking, head nodding, and jumping three times, and (3b) using large walking bouts to reorient the sensor data. (4) Turns are detected separately for head (4a) and trunk (4b) using gyroscope and accelerometer signals. (5) Head-on-body turns are identified by the diffference between head and trunk angular velocity, with thresholds (e.g., 5°/s) applied to detect significant independent head-on-body turn events.

Turns were identified when the angular velocity exceeded 15°/s, a threshold informed by literature on whole-body turning, which typically reports this value as the lower bound for meaningful rotational movements [62, 63]. To identify the start and end of a turn, a less aggressive FIR filter with an impulse duration of 0.383s was used to balance over-smoothing while ensuring accurate turn duration without misidentifying brief pauses as turn endings [60]. The start and stop points of a turn were identified when the rotational rate dropped below 5°/s to account for slowing and continuation of turns. The angular velocity curve between these points was then integrated to determine the amplitude of the turn.

To address instances where turns were executed in a jerky, non-smooth manner, we implemented an iterative process to consolidate turns in the same direction that occurred within 1/3 second of each other [62]. This specific time window was chosen based on the typical duration of brief pauses or hesitations during turning, ensuring that closely spaced movements were not mistakenly classified as separate events. Turns with an amplitude of less than 10 degrees were further examined for proximity to other nearby turns within the 1/3 second window. If such turns were identified, they were merged into a single turn to capture the full extent of the rotational movement. This process was iterated until no further nearby turns were identified in the same direction or until the cumulative duration of the merged turn exceeded 10 seconds. This approach prevents the exclusion of potential small turns caused by brief slowdowns or hesitations. Finally, small turns below 10 degrees in amplitude were discarded. This threshold was lower than the typical 40-degree threshold commonly employed in clinical studies [50, 51, 57, 61]. A threshold of 40 degrees is often used to avoid misclassifying curved walking, but we considered a smaller threshold here given the aim to generate normative data and to capture smaller angle turns that may be common for head movements. Identical procedures were used to detect thoracic and lumbar turns.

Head turns with respect to the body reference frame (head-on-body turns) were calculated using the same procedure as described above, but applied to the difference between the yaw angular velocity of the head and lumbar sensors after all sensors were aligned with gravity. This approach allowed us to distinguish head-in-space movements from head-on-body movements. We further classified head-on-body movements into volitional or stabilizing head-on-body turns by comparing the peak head-in-space and body-in-space angular velocities for each head-on-body turn. Volitional head-on-body turns were defined when the peak magnitude of the head-in-space angular velocity was greater than the peak magnitude of the lumbar-in-space angular velocity (i.e., the head was rotating faster than the body). Stabilizing head turns were defined when the peak magnitude of the head-in-space angular velocity was less than the peak magnitude of the lumbar-in-space angular velocity (i.e., the head was rotating slower than the body).

### G. Macro-Level Measures of Activity

#### 1) Step Count

To detect macro-level measures of mobility such as step count, the vertical acceleration signal from the lumbar sensor was smoothed by integrating and differentiating the signal using a Gaussian continuous wavelet transform. The initial contact of the foot was then identified as the minimum of the smoothed signal [64, 65]. If the number of steps in a bout was less than five, those steps were considered shuffling or standing and did not contribute to the step count.

#### 2) Wear Time

Activity counts were generated by summing the measured tri-axial acceleration over a one-second epoch [66]. These 1-second epochs were then aggregated to provide counts per minute, which represented the total intensity of movement within each minute. Extended periods of inactivity or zero counts were used to identify periods when the device was not being worn. Specifically, we considered the device to be unworn when we observed at least 90 consecutive minutes of zero counts. This method allowed for short bursts of activity (up to 2 minutes) within this period, which might be caused by incidental movement of an unworn device, without breaking the non-wear classification [67]. Wear time was calculated for each day, which was defined as the 24-hour period from midnight to midnight.

### H. Data Analysis

We analyzed wear time and compliance rates for head sensors over a 7-day period using descriptive statistics. To ensure consistent sensor orientation, we compared two alignment methods—calibration movements and long walking bouts—using paired t-tests across all turning metrics. Additionally, we established normative descriptive statistics, including the mean, range, median, and coefficient of variation (CV), for head-turning kinematics in healthy adults. We further explored the variance in head motion (as the dependent variable) explained by other continuous monitoring metrics, such as step count (as the independent variable), using linear mixed-effects models. These models accounted for repeated measures by including a random intercept for each participant. The strength of the relationships was assessed using the model’s adjusted R^2^ values [68]. Lastly, we identified the recommended wear time necessary to obtain reliable measures of head-turning kinematics.

To determine the optimal duration for reliable head turn data during daily life, we used intra-class correlation coefficients (ICC) with a two-way mixed-effects model (ICC [2,k]) to assess measurement reliability across multiple raters [69]. This statistical method assessed the consistency and reliability of turning characteristics, including turn peak velocity, turn angle amplitude, and number of turns across multiple days of wear. Analyses were restricted to days with a minimum of 10 hours of wear time to ensure sufficient data capture [70]. Specifically, ICC values below 0.50 were classified as indicating poor reliability, values between 0.50 and 0.75 as moderate reliability, values between 0.75 and 0.90 as good reliability, and values above 0.90 as excellent reliability [71]. We adopted 0.75 as a practical threshold for “good reliability,” consistent with common practices in gait analysis studies [55]. This approach aligns with common practices in gait analysis using IMUs, where two to three days of continuous monitoring is typically recommended for capturing reliable measures (ICC good reliability > 0.75) in movement patterns [55]. The relationship between independent head turns and step count was further examined to understand their potential association.

## III. Results

### A. Participant Compliance

Participants in the study (n = 24, 15 female) wore the sensors for an average of 16.38 hours per day (SD = 4.43). Each day required a minimum of 10 hours or wear to be considered as a full day [70]. All participants completed at least three consecutive days of wear, with an average of 6 days (SD = 1, min = 3, max = 9) of wear per participant. Five participants withdrew from the study before completing the 7 consecutive days of recording, but their data are included in the descriptive statistics. *Figure 2* provides an example of a single subject’s week of head turns per day.

**Fig. 2.**
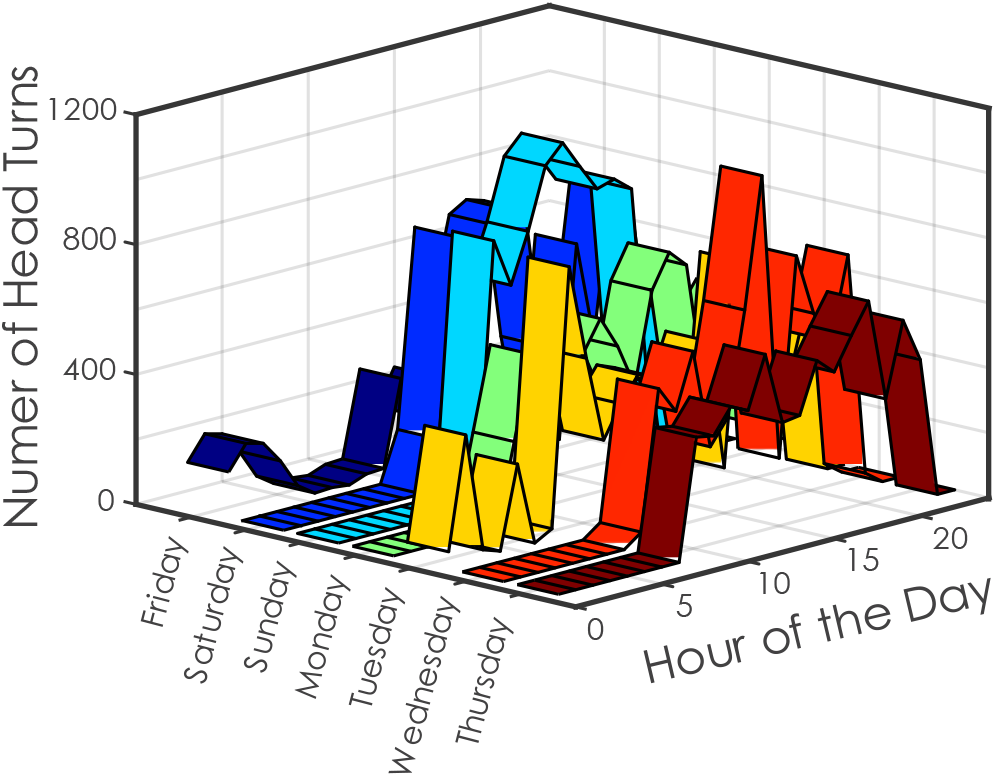
Representative data from a single subject showing the number of head turns per hour over a 24-hour period during a full week. These data illustrate temporal variations in head turn activity within a day and across different days of the week.

### B. Calibration Data

There was no significant difference between calibration and walking realignment methods for angular velocity (mean difference = 0.52°/s, 95% CI [-0.51, 1.54], p = 0.322) or number of head turns (mean difference = −0.61°/s, 95% CI [-11.13, 9.91], p = 0.910). However, head turn amplitude was significantly smaller in walking-based orientations compared to calibration-based orientations (mean difference −2.50°/s, 95% CI [-3.19, −1.82], p < 0.001).

### C. Descriptive Statistics of Normative Head and Lumbar Turns

Individuals completed an average (SD) of 5163 (1466) head turns and 2570 (746) lumbar turns per day (Table 1). The mean amplitude of turns was similar between head (58.18°, SD = 4.26°) and lumbar (60.97°, SD = 5.96°) regions. However, the mean peak velocity was greater for head turns (104.49°/s, SD = 12.08°/s) compared to lumbar turns (74.21°/s, SD = 6.07°/s). The variability in peak velocity was relatively high across all regions, with a CV of 0.83 (SD = 0.04) for the head and 0.83 (SD = 0.05) for the lumbar region (Table 2). However, when isolating small (<45°) and large head turns, the variability decreased (Table 2). The majority of head turn peak velocities and amplitudes during free-living daily life occurred at speeds less than 110°/s and with amplitudes smaller than 80° (Figure 3).

**TABLE I:**
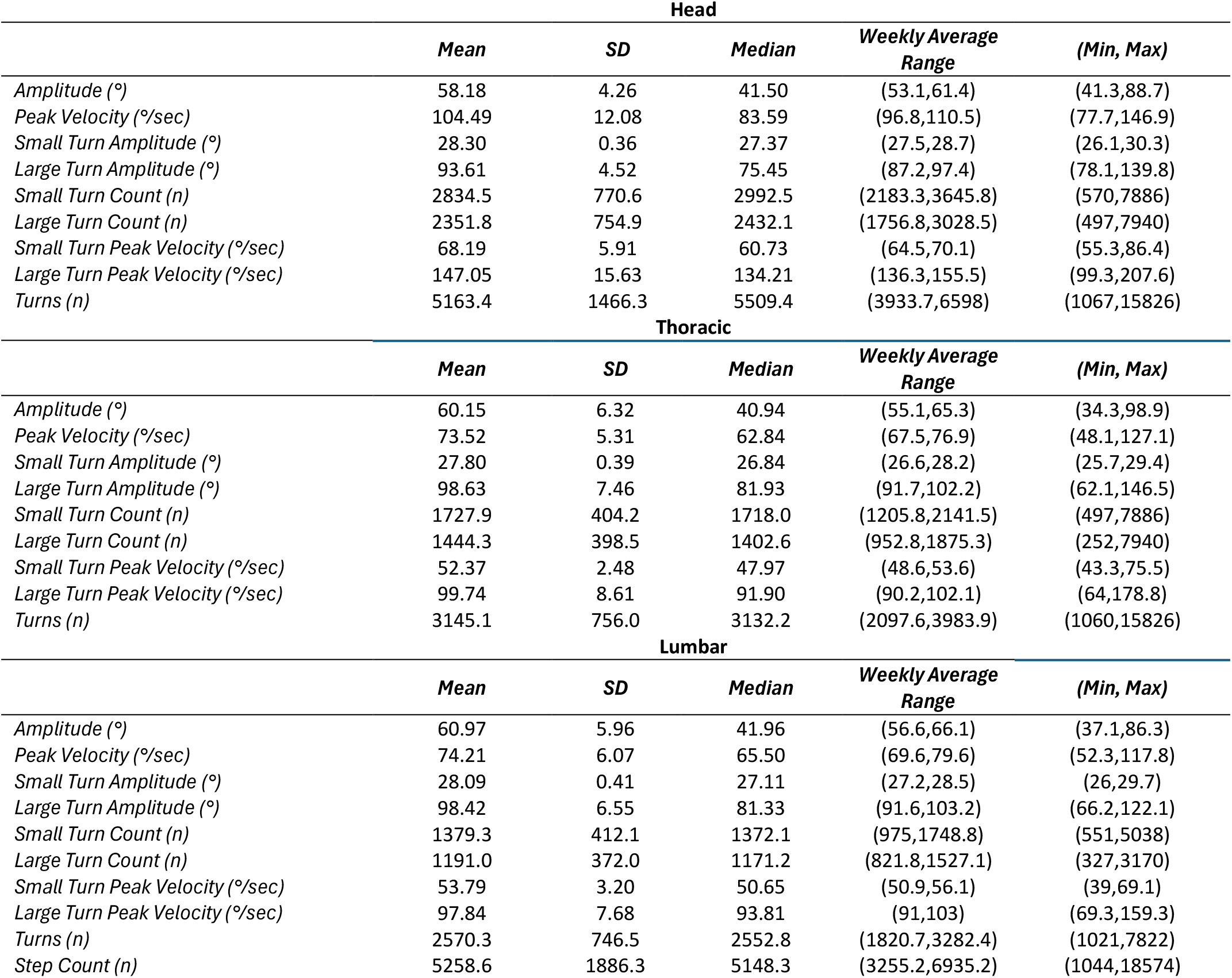
Summary STATISTICS (MEAN, STANDARD DEVIATION, MEDIAN, AND RANGE) FOR AMPLITUDE, PEAK VELOCITY, SMALL AND LARGE TURN AMPLITUDE AND VELOCITY, TURN COUNT FOR EACH BODY SEGMENT (Head, Thoracic, AND Lumbar), AND STEP COUNT. All MEAN, SD, MEDIAN, AND RANGE VALUES ARE BASED ON THE WEEKLY AVERAGE FOR EACH PARTICIPANT, WHILE THE INDIVIDUAL DAY MINIMUM AND MAXIMUM VALUES (Min, Max) ARE BASED ON SINGLE-DAY RECORDINGS.

| Head |  |  |  |  |  |
| --- | --- | --- | --- | --- | --- |
|  | Mean | SD | Median | Weekly Average Range | (Min, Max) |
| Amplitude (°) | 58.18 | 4.26 | 41.50 | (53.1,61.4) | (41.3,88.7) |
| Peak Velocity (°/sec) | 104.49 | 12.08 | 83.59 | (96.8,110.5) | (77.7,146.9) |
| Small Turn Amplitude (°) | 28.30 | 0.36 | 27.37 | (27.5,28.7) | (26.1,30.3) |
| Large Turn Amplitude (°) | 93.61 | 4.52 | 75.45 | (87.2,97.4) | (78.1,139.8) |
| Small Turn Count (n) | 2834.5 | 770.6 | 2992.5 | (2183.3,3645.8) | (570,7886) |
| Large Turn Count (n) | 2351.8 | 754.9 | 2432.1 | (1756.8,3028.5) | (497,7940) |
| Small Turn Peak Velocity (°/sec) | 68.19 | 5.91 | 60.73 | (64.5,70.1) | (55.3,86.4) |
| Large Turn Peak Velocity (°/sec) | 147.05 | 15.63 | 134.21 | (136.3,155.5) | (99.3,207.6) |
| Turns (n) | 5163.4 | 1466.3 | 5509.4 | (3933.7,6598) | (1067,15826) |
| Thoracic |  |  |  |  |  |
|  | Mean | SD | Median | Weekly Average Range | (Min, Max) |
| Amplitude (°) | 60.15 | 6.32 | 40.94 | (55.1,65.3) | (34.3,98.9) |
| Peak Velocity (°/sec) | 73.52 | 5.31 | 62.84 | (67.5,76.9) | (48.1,127.1) |
| Small Turn Amplitude (°) | 27.80 | 0.39 | 26.84 | (26.6,28.2) | (25.7,29.4) |
| Large Turn Amplitude (°) | 98.63 | 7.46 | 81.93 | (91.7,102.2) | (62.1,146.5) |
| Small Turn Count (n) | 1727.9 | 404.2 | 1718.0 | (1205.8,2141.5) | (497,7886) |
| Large Turn Count (n) | 1444.3 | 398.5 | 1402.6 | (952.8,1875.3) | (252,7940) |
| Small Turn Peak Velocity (°/sec) | 52.37 | 2.48 | 47.97 | (48.6,53.6) | (43.3,75.5) |
| Large Turn Peak Velocity (°/sec) | 99.74 | 8.61 | 91.90 | (90.2,102.1) | (64,178.8) |
| Turns (n) | 3145.1 | 756.0 | 3132.2 | (2097.6,3983.9) | (1060,15826) |
| Lumbar |  |  |  |  |  |
|  | Mean | SD | Median | Weekly Average Range | (Min, Max) |
| Amplitude (°) | 60.97 | 5.96 | 41.96 | (56.6,66.1) | (37.1,86.3) |
| Peak Velocity (°/sec) | 74.21 | 6.07 | 65.50 | (69.6,79.6) | (52.3,117.8) |
| Small Turn Amplitude (°) | 28.09 | 0.41 | 27.11 | (27.2,28.5) | (26,29.7) |
| Large Turn Amplitude (°) | 98.42 | 6.55 | 81.33 | (91.6,103.2) | (66.2,122.1) |
| Small Turn Count (n) | 1379.3 | 412.1 | 1372.1 | (975,1748.8) | (551,5038) |
| Large Turn Count (n) | 1191.0 | 372.0 | 1171.2 | (821.8,1527.1) | (327,3170) |
| Small Turn Peak Velocity (°/sec) | 53.79 | 3.20 | 50.65 | (50.9,56.1) | (39,69.1) |
| Large Turn Peak Velocity (°/sec) | 97.84 | 7.68 | 93.81 | (91,103) | (69.3,159.3) |
| Turns (n) | 2570.3 | 746.5 | 2552.8 | (1820.7,3282.4) | (1021,7822) |
| Step Count (n) | 5258.6 | 1886.3 | 5148.3 | (3255.2,6935.2) | (1044,18574) |

**TABLE II:**
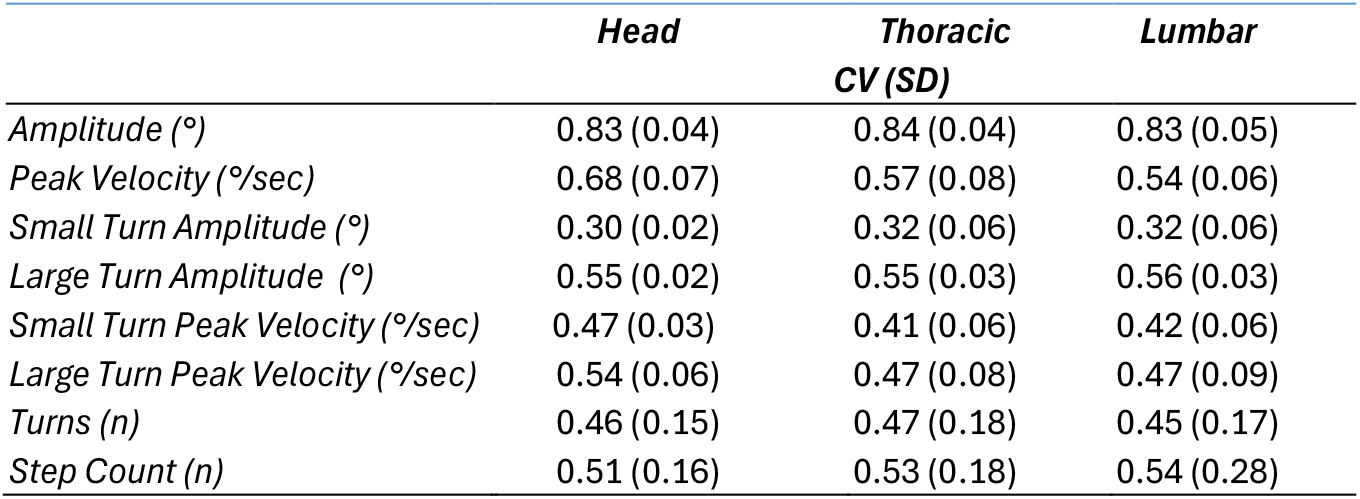
Coefficient OF VARIATION (CV) VALUES (MEAN AND STANDARD DEVIATION) FOR AMPLITUDE, PEAK VELOCITY, SMALL AND LARGE TURN AMPLITUDE, TURN COUNT, AND STEP COUNT ACROSS THREE BODY SEGMENTS (Head, Thoracic, AND Lumbar). These CV VALUES WERE CALCULATED FROM EACH INDIVIDUAL’S WEEKLY CV AND THEN AVERAGED ACROSS ALL SUBJECTS.

**Fig. 3.**
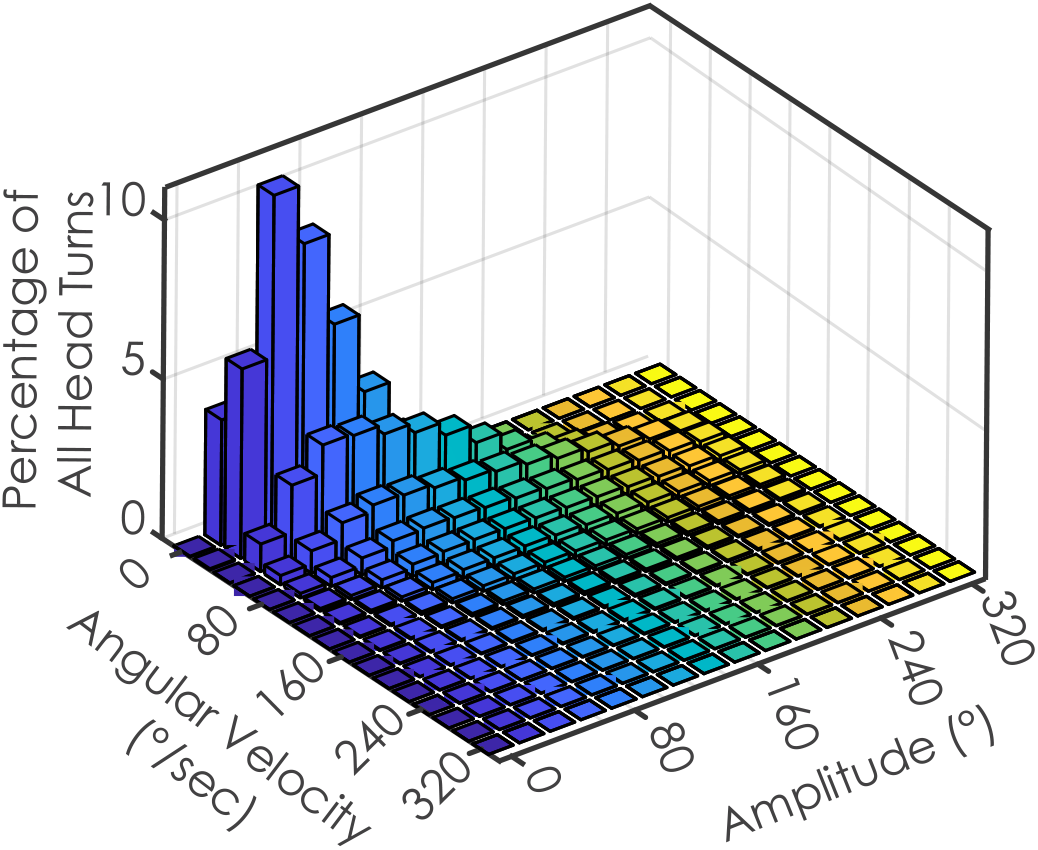
The distribution of the frequency of head turns across all participants, binned by speed and amplitude, during free-living daily life.

Participants completed an average of 4083 (SD = 1330) head turns independent from the trunk, with over 72% of all head turns occurring independently of trunk movement. These head-on-body turns were categorized as either stabilizing or volitional head turns, reflecting distinct patterns of segmental coordination (Table 3). Comparing kinematics of these turns with respect to the global reference frame (i.e. head-in-space amplitude and angular velocity), stabilizing turns were smaller and slower, with an average amplitude of 38.39° (SD = 7.68°) and peak velocity of 58.49°/s (SD = 11.32°/s), while volitional turns were characterized by larger amplitudes (88.53°, SD = 9.96°) and faster peak velocities (139.85°/s, SD = 27.41°/s).

**Table III.** Descriptive MEAN AND STANDARD DEVIATION (SD) VALUES FOR HEAD TURN AMPLITUDE AND VELOCITY, GIVEN FOR BODY-FIXED AND GLOBAL (I.E., SPACE) REFERENCE FRAMES, FOR STABILIZING (I.E., THE HEAD WAS ROTATING SLOWER THAN THE BODY) AND VOLITIONAL (I.E., THE HEAD WAS ROTATING FASTER THAN THE BODY) TURNS.

|  | *Stabilizing* | *Volitional* |
| --- | --- | --- |
|  | *Mean (SD)* |  |
| *Head-on-Body Turn Amplitude (°)* | 57.06 (9.44) | 61.95 (7.47) |
| *Head-on-Body Turn Angular Velocity (°/sec)* | 36.62 (4.9) | 38.12 (4.03) |
| *Head-in-Space Turn Amplitude (°)* | 38.39 (7.68) | 88.53 (9.96) |
| *Head-in-Space Turn Angular Velocity (°/sec)* | 58.49 (11.32) | 139.85 (27.41) |
| *Head Turn Count (n)* | 1548 (520) | 3145 (933) |

However, stabilizing and volitional head turns had comparable kinematics with respect to the body (i.e., head-on-body amplitude and angular velocity). The average amplitude was 57.06° (SD = 9.44°) for stabilizing turns and 61.95° (SD = 7.47°) for volitional turns. Similarly, the peak angular velocities were 36.62°/s (SD = 4.90°) for stabilizing turns and 38.12°/s (SD = 4.03°) for volitional turns. Overall, stabilizing turns were less frequent, averaging 1,548 turns per day (SD = 520), compared to volitional turns, which occurred an average of 3,146 turns per day (SD = 933).

### D. Independence of Head Turns

Participants averaged 5,233 (SD = 2,540) steps per day, with a mean of 43.1 steps per walking bout (SD = 17.9) and 385 steps per hour (SD = 264). Linear mixed models revealed a moderate relationship between the number of independent head turns and step count (adjusted r^2^ = 0.47) across all participants

### E. Wear Time for Reliability

On day 1, the ICC values indicated moderate reliability for amplitude (ICC = 0.60), while angular velocity (ICC = 0.78), the number of head turns (ICC = 0.88), head-on-body turns (ICC = 0.86), and step count (ICC = 0.89) showed good reliability. Using a two-day average, all metrics achieved excellent reliability (ICC > 0.9) relative to the week’s average (Figure 4).

**Fig. 4.**
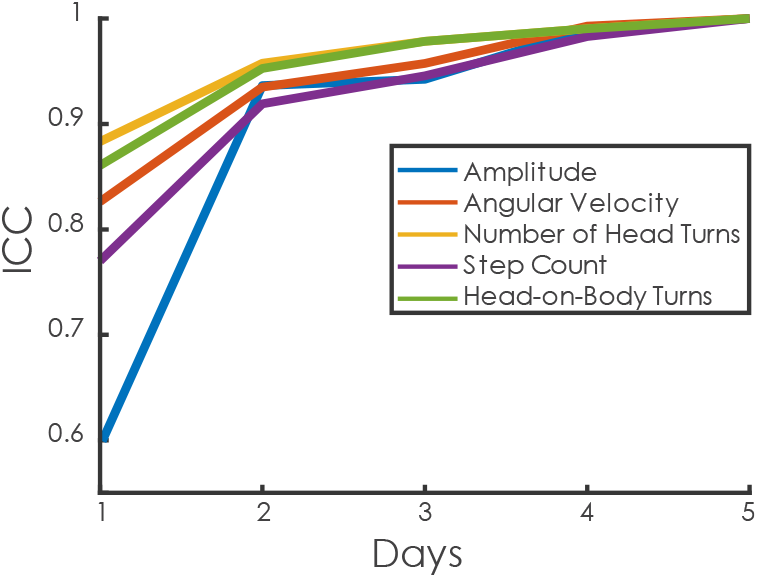
Intraclass correlation coefficients (ICC) values for amplitude, velocity, number of head turns, step count, and head-on-body turn based on the number of days included in the average compared to a 5-day average. All measures, except amplitude, showed good reliability (ICC > 0.75) on the first day. After day two, all measures exhibited excellent reliability (ICC > 0.9) with the 5-day average.

## IV. Discussion

The purpose of this study was to examine the feasibility of capturing head kinematics during free-living daily activities over a 7-day period and to establish normative data for young adults. Specifically, we sought to determine feasibility by answering key issues related to continuous monitoring of head movements during daily life: 1) participant compliance; 2) methods for achieving repeatable sensor orientation; 3) normative descriptive statistics; 4) relationships between head motion and other continuous monitoring metrics; and 5) recommendations for the amount of wear time needed to obtain reliable measures. We demonstrated that quantifying head kinematics, including the number, frequency, amplitude, and velocity of head turns, is feasible in free-living daily life based on high compliance and repeatable methods to realign sensor-data, advancing prior work in controlled laboratory or simulated community ambulation tasks [39, 48-51, 72, 73]. We also provided normative data on head turns, lumbar turns, and head-trunk coupling in healthy, young adults to complement previous studies on free-living whole body turns in patient populations [17, 57]. We showed that the number of head turns are not well explained by traditional measures of activity. Finally, our results suggest that reliable data on free-living head turns can be achieved in as little as two days. Combined, these findings lay the groundwork for quantifying head kinematics in future clinical applications.

### A. Participant Compliance

Patient compliance in our study of head turns was comparable to that reported in other continuous monitoring studies [70, 74-77]. We achieved a compliance of 82.8% over the 7-day monitoring period with participants wearing the three sensors for an average of 15.9 hours per day. The high compliance observed in our study can likely be attributed to three key factors: comfortable head sensor placement, easy once-a-day application, and long battery life such that the sensors did not require charging. Similar to the activPAL (PAL Technologies Ltd, Glasgow, Scotland), we used adhesive attachment rather than a belt system, allowing users to easily attach the IMUs and then proceed with their daily life [78, 79]. Furthermore, the IMUs did not require charging during the entire study duration, which reduced the burden on participants [80, 81].

### B. Head Turn Kinematics

Our findings on head turning kinematics during free-living activities revealed much smaller and slower head turns than those reported in healthy individuals by previous in-lab assessments [39, 50, 82-84]. During in-lab assessments, the prescribed clinical tasks are designed to elicit a full range of motion with little variability in the amplitude or speed of head turn. Conversely, individuals exhibit high variability in head movements throughout the day, and often make small changes in head orientation to navigate [47]. While, the inclusion of a lower threshold to define a head turn in our study may also contribute to the smaller turns, which, in turn, affect the overall descriptive statistics of head movement, skewing the results, our descriptive statistics align with head turn amplitude and speed during a simulated 15-minute community ambulatory walk [51]. These results suggest that obtaining ecologically valid measures of head kinematics may be obtained from free-living daily life or carefully designed simulated tasks. However, a 15-minute community ambulation task may be challenging to implement in clinical settings due to time and space limitations, while continuous monitoring during free-living daily life enables remote monitoring assessments without disrupting clinical workflows.

Our study observed that over 72 percent of the head turns occurred independently of the trunk during free-living daily life, highlighting the prevalence of isolated head movements in daily activities. Monitoring the frequency of head-on-body turns may be clinically useful; when walking and turning, people with Parkinson’s disease commonly use more head-with-body turns (i.e., ‘en-bloc’ turns) instead of sequential rotations of the head, then the body, due to rigidity and bradykinesia [85-88]. This frequency of head-on-body turns in daily life also underscores that movements throughout the day involve both head-with-body (i.e., synchronous head and body turns) and head-on-body (i.e., isolated head motion) turns. For example, seated activities like desk work or driving an automobile can require volitional head turns while the trunk remains stationary. Notably, head movements during sedentary periods, which can include non-verbal communication / gesture (i.e. head nods or shakes) may still reflect important aspects of an individual’s daily function and impact quality of life [89-93]. Based on the speed of these turns in space, volitional head turns serve to change gaze or head position, perhaps to obtain new visual information about the environment. Conversely, stabilizing head turns may maintain visual fixation or smooth pursuit of an object while the rest of the body is turning since the speed of the stabilizing head turns was slower than the maximum smooth pursuit velocity [94]. Based on these different functions of volitional and stabilizing head turns, monitoring these different head turns may provide insight into the effects of ocular or vestibular dysfunction, even during sedentary behaviors.

### C. Independence of Head Turns and Clinical Value

Despite similar mean counts, head turns were only moderately associated with step count, suggesting head kinematics can serve as a complementary measure of function that is largely unexplained by step count. Step count is frequently employed as a proxy measure for functional mobility and overall activity level [14, 95, 96], but step count fails to capture clinically-relevant micro-level aspects of mobility [1, 17, 56, 57, 97]. Head turn metrics may provide complementary information about the complexity and variability of daily movements, especially those involving the head and neck. For example, changes in head turn amplitude or velocity during daily activities can serve as early indicators of disease progression in people with Parkinson’s disease [56, 98], improvements in head turn frequency and velocity may signify successful rehabilitation in people with vestibular hypofunction [49-51], and comparing pre- and post-treatment head movement patterns could assess the effectiveness of interventions for neck pain or cervical spine injuries [84]. The number of head turns offers a promising metric for assessing functional mobility, providing insights that complement traditional measures like step count. The present results support incorporating head kinematics into broader assessments of daily mobility, combining both gross motor activities (e.g., walking) and finer movements (e.g., head turns) to create a more comprehensive understanding of movement patterns in daily life.

### D. Recommendation for Implementation

Our findings reveal that reliable measurements of head turns (e.g., count, speed, amplitude) can be obtained after just two, 10-hour days of recording, aligning with studies on gait and sway variability that demonstrate similar reliability within two to three days of wear [55, 77]. This high reliability suggests that representative data on head movements can be gathered without imposing prolonged monitoring periods on participants. To further reduce participant’s burden, we found that individual calibration movements were unnecessary, as walking bouts provided sufficient data for sensor reorientation and yielded comparable results to calibration-based results. Regarding sensor attachment, most participants tolerated both latex-free double-sided tape and Tegaderm well. However, we recommend changing the adhesive midweek to allow skin aeration. Some participants found Tegaderm irritating, and for those less prone to sweating, double-sided tape alone proved sufficient. These options offer flexibility in accommodating individual comfort and skin sensitivity.

### E. Limitations

While our study provides valuable insights into head turn kinematics, such as the number, speed, and amplitude, during daily life, several limitations should be acknowledged. Our sample was relatively small and homogeneous, primarily consisting of young, healthy adults, many of whom were desk workers. This limited diversity may restrict the generalizability of our findings to other age groups or more physically active individuals. Additionally, we lacked detailed information about specific daily activities that could influence head kinematics, such as cycling or skiing. Classifying activities may provide context for understanding factors that modulate head movements in real-world settings.

Future research should incorporate cross-sectional studies to compare head movement patterns between healthy and impaired populations, helping to identify mobility-related biomarkers in conditions such as acquired brain injury or Parkinson’s disease. Longitudinal studies are also essential to track changes over time, whether due to natural recovery, disease progression, or rehabilitation. Expanding sample diversity, incorporating detailed activity logs, and integrating these study designs will enhance our understanding of head movement kinematics in daily life and their implications for clinical assessments and interventions.

## V. Conclusion

This study demonstrates the feasibility of capturing head turn kinematics during free-living daily activities over an extended period, advancing previous work in controlled settings. Our findings provide normative data on head turns, thoracic turns, lumbar turns, and head-trunk coupling in healthy, young adults. Key outcomes included high participant compliance, establishment of normative data for head turn metrics, and identification of independent head turns. The relatively weak relationship between head turns and step count emphasizes the need for comprehensive assessments of daily movement patterns beyond just the accumulation of steps. Our study showcases the potential of wearable sensors in capturing head kinematics during daily life, highlighting its importance in assessing function. We recommend a minimum of two days of wear, with at least 10 hours of data collection daily, and suggest using large walking bouts for sensory re-orientation, reducing patient burden while maintaining accuracy in real-world settings.

## Acknowledgment

Thank you to Noemi Montemurro, Christina Geisler, and Corinne Mayfield for assisting patient recruitment, data collection, and data processing.

## References

[1] L. Rochester, S. F. M. Chastin, S. Lord, K. Baker, and D. J. Burn, “Understanding the impact of deep brain stimulation on ambulatory activity in advanced Parkinson’s disease,” (in English), J Neurol, vol. 259, no. 6, pp. 1081–1086, Jun 2012, doi: 10.1007/s00415-011-6301-9.

[2] J. T. Cavanaugh, T. D. Ellis, G. M. Earhart, M. P. Ford, K. B. Foreman, and L. E. Dibble, “Capturing Ambulatory Activity Decline in Parkinson’s Disease,” (in English), J Neurol Phys Ther, vol. 36, no. 2, pp. 51–57, Jun 2012, doi: 10.1097/NPT.0b013e318254ba7a.

[3] M. E. de Leeuwerk, P. Bor, H. P. van der Ploeg, V. de Groot, M. van der Schaaf, and M. van der Leeden, “The effectiveness of physical activity interventions using activity trackers during or after inpatient care: a systematic review and meta-analysis of randomized controlled trials,” Int J Behav Nutr Phy, vol. 19, no. 1, p. 59, 2022.

[4] K. Szeto, J. Arnold, B. Singh, B. Gower, C. E. Simpson, and C. Maher, “Interventions using wearable activity trackers to improve patient physical activity and other outcomes in adults who are hospitalized: a systematic review and meta-analysis,” JAMA Network Open, vol. 6, no. 6, pp. e2318478–e2318478, 2023.

[5] K. Taraldsen, S. F. M. Chastin, I. I. Riphagen, B. Vereijken, and J. L. Helbostad, “Physical activity monitoring by use of accelerometer-based body-worn sensors in older adults: A systematic literature review of current knowledge and applications,” (in English), Maturitas, vol. 71, no. 1, pp. 13–19, Jan 2012, doi: 10.1016/j.maturitas.2011.11.003.

[6] E. D. de Bruin, A. Hartmann, D. Uebelhart, K. Murer, and W. Zijlstra, “Wearable systems for monitoring mobility-related activities in older people: a systematic review,” (in English), Clin Rehabil, vol. 22, no. 10-11, pp. 878–895, 2008, doi: 10.1177/0269215508090675.

[7] R. I. Spain et al., “Body-worn motion sensors detect balance and gait deficits in people with multiple sclerosis who have normal walking speed,” (in English), Gait Posture, vol. 35, no. 4, pp. 573–578, Apr 2012, doi: 10.1016/j.gaitpost.2011.11.026.

[8] R. I. Spain, M. Mancini, F. B. Horak, and D. Bourdette, “Body-worn sensors capture variability, but not decline, of gait and balance measures in multiple sclerosis over 18 months,” (in English), Gait Posture, vol. 39, no. 3, pp. 958–964, Mar 2014, doi: 10.1016/j.gaitpost.2013.12.010.

[9] M. Mancini, P. Carlson-Kuhta, C. Zampieri, J. G. Nutt, L. Chiari, and F. B. Horak, “Postural sway as a marker of progression in Parkinson’s disease: A pilot longitudinal study,” (in English), Gait Posture, vol. 36, no. 3, pp. 471–476, Jul 2012, doi: 10.1016/j.gaitpost.2012.04.010.

[10] J. M. Hausdorff, H. K. Edelberg, S. L. Mitchell, A. L. Goldberg, and J. Y. Wei, “Increased gait unsteadiness in community-dwelling elderly fallers,” (in English), Arch Phys Med Rehab, vol. 78, no. 3, pp. 278–283, Mar 1997, doi: Doi 10.1016/S0003-9993(97)90034-4.

[11] G. A. Mbourou, Y. Lajoie, and N. Teasdale, “Step length variability at gait initiation in elderly fallers and non-fallers, and young adults,” (in English), Gerontology, vol. 49, no. 1, pp. 21–26, Jan-Feb 2003, doi: 10.1159/000066506.

[12] A. H. Newstead, J. G. Walden, and A. J. Gitter, “Gait variables differentiating fallers from nonfallers,” J Geriatr Phys Ther, vol. 30, no. 3, pp. 93–101, 2007, doi: 10.1519/00139143-200712000-00003.

[13] L. A. King et al., “Instrumenting the Balance Error Scoring System for Use With Patients Reporting Persistent Balance Problems After Mild Traumatic Brain Injury,” (in English), Arch Phys Med Rehab, vol. 95, no. 2, pp. 353–359, Feb 2014, doi: 10.1016/j.apmr.2013.10.015.

[14] K. S. Hall et al., “Systematic review of the prospective association of daily step counts with risk of mortality, cardiovascular disease, and dysglycemia,” (in English), Int J Behav Nutr Phy, vol. 17, no. 1, Jun 20 2020, doi: ARTN 78 10.1186/s12966-020-00978-9.

[15] S. Del Din et al., “Analysis of Free-Living Gait in Older Adults With and Without Parkinson’s Disease and With and Without a History of Falls: Identifying Generic and Disease-Specific Characteristics,” The Journals of Gerontology: Series A, vol. 74, no. 4, pp. 500–506, 2017, doi: 10.1093/gerona/glx254.

[16] M. El-Gohary et al., “Continuous Monitoring of Turning in Patients with Movement Disability,” (in English), Sensors-Basel, vol. 14, no. 1, pp. 356–369, Jan 2014, doi: 10.3390/s140100356.

[17] M. Mancini et al., “Continuous monitoring of turning in Parkinson’s disease: Rehabilitation potential,” (in English), Neurorehabilitation, vol. 37, no. 1, pp. 3–10, 2015, doi: 10.3233/Nre-151236.

[18] L. Baroudi, K. Barton, S. M. Cain, and K. A. Shorter, “Classification of human walking context using a single-point accelerometer,” Scientific Reports, vol. 14, no. 1, p. 3039, 2024/02/06 2024, doi: 10.1038/s41598-024-53143-8.

[19] L. Baroudi, S. M. Cain, K. A. Shorter, and K. Barton, “Enhancing the Efficacy of Lower-body Assistive Devices Through the Understanding of Human Movement in the Real World,” in 2023 IEEE International Conference on Robotics and Automation (ICRA), 29 May-2 June 2023 2023, pp. 11351–11358, doi: 10.1109/ICRA48891.2023.10161051.

[20] A. Cereatti et al., “ISB recommendations on the definition, estimation, and reporting of joint kinematics in human motion analysis applications using wearable inertial measurement technology,” Journal of Biomechanics, vol. 173, p. 112225, 2024/08/01/ 2024, doi: 10.1016/j.jbiomech.2024.112225.

[21] S. Del Din, A. Godfrey, C. Mazzà, S. Lord, and L. Rochester, “Free-living monitoring of Parkinson’s disease: Lessons from the field,” Movement Disorders, vol. 31, no. 9, pp. 1293–1313, 2016, doi: 10.1002/mds.26718.

[22] S. D. Din, A. Godfrey, and L. Rochester, “Validation of an Accelerometer to Quantify a Comprehensive Battery of Gait Characteristics in Healthy Older Adults and Parkinson’s Disease: Toward Clinical and at Home Use,” IEEE Journal of Biomedical and Health Informatics, vol. 20, no. 3, pp. 838–847, 2016, doi: 10.1109/JBHI.2015.2419317.

[23] V. V. Shah et al., “Does gait bout definition influence the ability to discriminate gait quality between people with and without multiple sclerosis during daily life?,” Gait Posture, vol. 84, pp. 108–113, 2021/02/01/ 2021, doi: 10.1016/j.gaitpost.2020.11.024.

[24] V. V. Shah et al., “Effect of Bout Length on Gait Measures in People with and without Parkinson’s Disease during Daily Life,” Sensors-Basel, vol. 20, no. 20, p. 5769, 2020. [Online]. Available: https://www.mdpi.com/1424-8220/20/20/5769.

[25] V. V. Shah et al., “Digital Biomarkers of Mobility in Parkinson’s Disease During Daily Living,” Journal of Parkinson’s Disease, vol. 10, pp. 1099–1111, 2020, doi: 10.3233/JPD-201914.

[26] V. V. Shah et al., “Laboratory versus daily life gait characteristics in patients with multiple sclerosis, Parkinson’s disease, and matched controls,” Journal of NeuroEngineering and Rehabilitation, vol. 17, no. 1, p. 159, 2020/12/01 2020, doi: 10.1186/s12984-020-00781-4.

[27] M. D. F. R. W. Baloh, Baloh and Honrubia’s Clinical Neurophysiology of the Vestibular System. United Kingdom: United Kingdom: Oxford University Press, 2010.

[28] R. L. Cromwell, R. A. Newton, and L. G. Carlton, “Horizontal plane head stabilization during locomotor tasks,” (in English), J Motor Behav, vol. 33, no. 1, pp. 49–58, Mar 2001, doi: Doi 10.1080/00222890109601902.

[29] J. M. Goldberg and K. E. Cullen, “Vestibular control of the head: possible functions of the vestibulocollic reflex,” (in English), Experimental Brain Research, vol. 210, no. 3-4, pp. 331–345, May 2011, doi: 10.1007/s00221-011-2611-5.

[30] R. Grasso, S. Glasauer, Y. Takei, and A. Berthoz, “The predictive brain: Anticipatory control of head direction for the steering of locomotion,” (in English), Neuroreport, vol. 7, no. 6, pp. 1170–1174, Apr 26 1996, doi: Doi 10.1097/00001756-199604260-00015.

[31] R. Grasso, P. Prévost, Y. P. Ivanenko, and A. Berthoz, “Eye-head coordination for the steering of locomotion in humans:: an anticipatory synergy,” (in English), Neurosci Lett, vol. 253, no. 2, pp. 115–118, Sep 4 1998, doi: Doi 10.1016/S0304-3940(98)00625-9.

[32] P. Hausamann, M. Daumer, P. R. MacNeilage, and S. Glasauer, “Ecological Momentary Assessment of Head Motion: Toward Normative Data of Head Stabilization,” (in English), Front Hum Neurosci, vol. 13, Jun 4 2019, doi: ARTN 179 10.3389/fnhum.2019.00179.

[33] E. Hirasaki, S. T. Moore, T. Raphan, and B. Cohen, “Effects of walking velocity on vertical head and body movements during locomotion,” (in English), Experimental Brain Research, vol. 127, no. 2, pp. 117–130, Jul 1999, doi: DOI 10.1007/s002210050781.

[34] T. Mergner, C. Siebold, G. Schweigart, and W. Becker, “Human Perception of Horizontal Trunk and Head Rotation in Space during Vestibular and Neck Stimulation,” (in English), Experimental Brain Research, vol. 85, no. 2, pp. 389–404, 1991. [Online]. Available: <Go to ISI>://WOS:A1991FU18600013.

[35] J. O. Nnodim and R. L. Yung, “Balance and its Clinical Assessment in Older Adults - A Review,” J Geriatr Med Gerontol, vol. 1, no. 1, 2015, doi: 10.23937/2469-5858/1510003.

[36] T. Pozzo, A. Berthoz, and L. Lefort, “Head Stabilization during Various Locomotor Tasks in Humans. 1. Normal Subjects,” (in English), Experimental Brain Research, vol. 82, no. 1, pp. 97–106, 1990. [Online]. Available: <Go to ISI>://WOS:A1990EA64000011.

[37] T. Pozzo, A. Berthoz, L. Lefort, and E. Vitte, “Head Stabilization during Various Locomotor Tasks in Humans. 2. Patients with Bilateral Peripheral Vestibular Deficits,” (in English), Experimental Brain Research, vol. 85, no. 1, pp. 208–217, 1991. [Online]. Available: <Go to ISI>://WOS:A1991FN93200022.

[38] M. Saglam, S. Glasauer, and N. Lehnen, “Vestibular and cerebellar contribution to gaze optimality,” (in English), Brain, vol. 137, pp. 1080–1094, Apr 2014, doi: 10.1093/brain/awu006.

[39] D. Solomon, V. Kumar, R. A. Jenkins, and J. Jewell, “Head control strategies during whole-body turns,” (in English), Experimental Brain Research, vol. 173, no. 3, pp. 475–486, Aug 2006, doi: 10.1007/s00221-006-0393-y.

[40] S. T. Moore, E. Hirasaki, T. Raphan, and B. Cohen, “The Human Vestibulo-Ocular Reflex during Linear Locomotion,” Annals of the New York Academy of Sciences, vol. 942, no. 1, pp. 139–147, 2001, doi: 10.1111/j.1749-6632.2001.tb03741.x.

[41] K. Berg, S. Wood-Dauphinee, and J. Williams, “The Balance Scale: reliability assessment with elderly residents and patients with an acute stroke,” Scandinavian journal of rehabilitation medicine, vol. 27, no. 1, pp. 27–36, 1995.

[42] A. S. Cook and M. H. Woollacott, Motor control: theory and practical applications. Lippincott Williams and Wilkins, 2001.

[43] F. Franchignoni, F. Horak, M. Godi, A. Nardone, and A. Giordano, “Using psychometric techniques to improve the Balance Evaluation System’s Test: the mini-BESTest,” Journal of rehabilitation medicine: official journal of the UEMS European Board of Physical and Rehabilitation Medicine, vol. 42, no. 4, p. 323, 2010.

[44] F. B. Horak, D. M. Wrisley, and J. Frank, “The balance evaluation systems test (BESTest) to differentiate balance deficits,” Phys Ther, vol. 89, no. 5, pp. 484–498, 2009.

[45] A. L. Leddy, B. E. Crowner, and G. M. Earhart, “Functional Gait Assessment and Balance Evaluation System Test: Reliability, Validity, Sensitivity, and Specificity for Identifying Individuals With Parkinson Disease Who Fall,” Phys Ther, vol. 91, no. 1, pp. 102–113, 2011, doi: 10.2522/ptj.20100113.

[46] S. Whitney, M. Hudak, and G. Marchetti, “The dynamic gait index relates to self-reported fall history in individuals with vestibular dysfunction,” Journal of Vestibular Research, vol. 10, no. 2, pp. 99–105, 2000.

[47] D. M. Wrisley, G. F. Marchetti, D. K. Kuharsky, and S. L. Whitney, “Reliability, internal consistency, and validity of data obtained with the functional gait assessment,” Phys Ther, vol. 84, no. 10, pp. 906–918, 2004.

[48] J. L. Millar, O. A. Zobeiri, W. H. Souza, M. C. Schubert, and K. E. Cullen, “Head movement kinematics are differentially altered for extended versus short duration gait exercises in individuals with vestibular loss,” Scientific Reports, vol. 13, no. 1, p. 16213, 2023/09/27 2023, doi: 10.1038/s41598-023-42441-2.

[49] O. A. Zobeiri, G. M. Mischler, S. A. King, R. F. Lewis, and K. E. Cullen, “Effects of vestibular neurectomy and neural compensation on head movements in patients undergoing vestibular schwannoma resection,” Scientific Reports, vol. 11, no. 1, p. 517, 2021/01/12 2021, doi: 10.1038/s41598-020-79756-3.

[50] S. S. Paul, L. E. Dibble, R. G. Walther, C. Shelton, R. K. Gurgel, and M. E. Lester, “Characterization of Head-Trunk Coordination Deficits After Unilateral Vestibular Hypofunction Using Wearable Sensors,” (in English), Jama Otolaryngol, vol. 143, no. 10, pp. 1008–1014, Oct 2017, doi: 10.1001/jamaoto.2017.1443.

[51] S. S. Paul, L. E. Dibble, R. G. Walther, C. Shelton, R. K. Gurgel, and M. E. Lester, “Reduced Purposeful Head Movements During Community Ambulation Following Unilateral Vestibular Loss,” (in English), Neurorehabilitation and Neural Repair, vol. 32, no. 4-5, pp. 309–316, Apr-May 2018, doi: 10.1177/1545968318770271.

[52] O. A. Zobeiri, L. Wang, J. L. Millar, M. C. Schubert, and K. E. Cullen, “Head movement kinematics are altered during balance stability exercises in individuals with vestibular schwannoma,” Journal of NeuroEngineering and Rehabilitation, vol. 19, no. 1, p. 120, 2022.

[53] R. Aryan, O. A. Zobeiri, J. L. Millar, M. C. Schubert, and K. E. Cullen, “Effect of vestibular loss on head-on-trunk stability in individuals with vestibular schwannoma,” Scientific reports, vol. 14, no. 1, p. 3512, 2024.

[54] L. Wang, O. A. Zobeiri, J. L. Millar, W. Souza Silva, M. C. Schubert, and K. E. Cullen, “Continuous head motion is a greater motor control challenge than transient head motion in patients with loss of vestibular function,” Neurorehabilitation and neural repair, vol. 35, no. 10, pp. 890–902, 2021.

[55] B. M. Meyer et al., “How Much Data Is Enough? A Reliable Methodology to Examine Long-Term Wearable Data Acquisition in Gait and Postural Sway,” Sensors-Basel, vol. 22, no. 18, p. 6982, 2022. [Online]. Available: https://www.mdpi.com/1424-8220/22/18/6982.

[56] M. Mancini et al., “Continuous Monitoring of Turning Mobility and Its Association to Falls and Cognitive Function: A Pilot Study,” The Journals of Gerontology: Series A, vol. 71, no. 8, pp. 1102–1108, 2016, doi: 10.1093/gerona/glw019.

[57] S. Stuart et al., “Analysis of Free-Living Mobility in People with Mild Traumatic Brain Injury and Healthy Controls: Quality over Quantity,” J Neurotrauma, vol. 37, no. 1, pp. 139–145, Jan 1 2020, doi: 10.1089/neu.2019.6450.

[58] R. Moe-Nilssen, “A new method for evaluating motor control in gait under real-life environmental conditions. Part 1: The instrument,” Clinical Biomechanics, vol. 13, no. 4, pp. 320–327, 1998/06/01/ 1998, doi: 10.1016/S0268-0033(98)00089-8.

[59] R. Moe-Nilssen and J. L. Helbostad, “Trunk accelerometry as a measure of balance control during quiet standing,” Gait Posture, vol. 16, no. 1, pp. 60–68, 2002/08/01/ 2002, doi: 10.1016/S0966-6362(01)00200-4.

[60] V. V. Shah et al., “Inertial Sensor Algorithms to Characterize Turning in Neurological Patients With Turn Hesitations,” (in English), Ieee T Bio-Med Eng, vol. 68, no. 9, pp. 2615–2625, Sep 2021, doi: 10.1109/Tbme.2020.3037820.

[61] V. V. Shah et al., “Inertial Sensor Algorithms to Characterize Turning in Neurological Patients With Turn Hesitations,” (in eng), IEEE Trans Biomed Eng, vol. 68, no. 9, pp. 2615–2625, Sep 2021, doi: 10.1109/tbme.2020.3037820.

[62] L. Parrington, D. A. Jehu, P. C. Fino, S. Pearson, M. El-Gohary, and L. A. King, “Validation of an inertial sensor algorithm to quantify head and trunk movement in healthy young adults and individuals with mild traumatic brain injury,” Sensors-Basel, vol. 18, no. 12, p. 4501, 2018.

[63] S. Pearson, M. Mancini, M. El-Gohary, J. McNames, and F. Horak, “Turn detection and characterization with inertial sensors,” in International Congress on Sports Science Research and Technology Support, 2013, vol. 2: SciTePress, pp. 19–22.

[64] J. McCamley, M. Donati, E. Grimpampi, and C. Mazzà, “An enhanced estimate of initial contact and final contact instants of time using lower trunk inertial sensor data,” (in eng), Gait Posture, vol. 36, no. 2, pp. 316–8, Jun 2012, doi: 10.1016/j.gaitpost.2012.02.019.

[65] A. Paraschiv-Ionescu, C. J. Newman, L. Carcreff, C. N. Gerber, S. Armand, and K. Aminian, “Locomotion and cadence detection using a single trunk-fixed accelerometer: validity for children with cerebral palsy in daily life-like conditions,” Journal of NeuroEngineering and Rehabilitation, vol. 16, no. 1, p. 24, 2019/02/04 2019, doi: 10.1186/s12984-019-0494-z.

[66] Actigraph, “Actigraph GT1M Monitor/ActiTrainer and ActiLife Lifestyle Monitor software user manual,” ed: Actigraph Pensacola, FL, 2007.

[67] L. Choi, Z. Liu, C. E. Matthews, and M. S. Buchowski, “Validation of accelerometer wear and nonwear time classification algorithm,” (in eng), Med Sci Sports Exerc, vol. 43, no. 2, pp. 357–64, Feb 2011, doi: 10.1249/MSS.0b013e3181ed61a3.

[68] P. Schober, C. Boer, and L. A. Schwarte, “Correlation coefficients: appropriate use and interpretation,” Anesthesia & analgesia, vol. 126, no. 5, pp. 1763–1768, 2018.

[69] C. Tudor-Locke, L. Burkett, J. P. Reis, B. E. Ainsworth, C. A. Macera, and D. K. Wilson, “How many days of pedometer monitoring predict weekly physical activity in adults?,” (in eng), Prev Med, vol. 40, no. 3, pp. 293–8, Mar 2005, doi: 10.1016/j.ypmed.2004.06.003.

[70] A. Chan, D. Chan, H. Lee, C. C. Ng, and A. H. L. Yeo, “Reporting adherence, validity and physical activity measures of wearable activity trackers in medical research: A systematic review,” International Journal of Medical Informatics, vol. 160, p. 104696, 2022/04/01/ 2022, doi: 10.1016/j.ijmedinf.2022.104696.

[71] T. K. Koo and M. Y. Li, “A Guideline of Selecting and Reporting Intraclass Correlation Coefficients for Reliability Research,” Journal of Chiropractic Medicine, vol. 15, no. 2, pp. 155–163, 2016/06/01/ 2016, doi: 10.1016/j.jcm.2016.02.012.

[72] T. Mijovic, J. Carriot, A. Zeitouni, and K. E. Cullen, “Head Movements in Patients with Vestibular Lesion: A Novel Approach to Functional Assessment in Daily Life Setting,” (in English), Otol Neurotol, vol. 35, no. 10, pp. E348–E357, Dec 2014. [Online]. Available: <Go to ISI>://WOS:000345157200015.

[73] T. Pozzo, A. Berthoz, and L. Lefort, “Head stabilization during various locomotor tasks in humans,” Experimental Brain Research, vol. 82, no. 1, pp. 97–106, 1990/08/01 1990, doi: 10.1007/BF00230842.

[74] C. Buckley et al., “Quantifying Reliable Walking Activity with a Wearable Device in Aged Residential Care: How Many Days Are Enough?,” (in eng), Sensors (Basel), vol. 20, no. 21, Nov 5 2020, doi: 10.3390/s20216314.

[75] J. E. Sasaki et al., “Number of days required for reliably estimating physical activity and sedentary behaviour from accelerometer data in older adults,” Journal of sports sciences, vol. 36, no. 14, pp. 1572–1577, 2018.

[76] B. Western et al., “How many days of continuous physical activity monitoring reliably represent time in different intensities in cancer survivors,” Plos One, vol. 18, no. 4, p. e0284881, 2023.

[77] K. S. van Schooten, S. M. Rispens, P. J. M. Elders, P. Lips, J. H. van Dieën, and M. Pijnappels, “Assessing Physical Activity in Older Adults: Required Days of Trunk Accelerometer Measurements for Reliable Estimation,” (in English), Journal of Aging and Physical Activity, vol. 23, no. 1, pp. 9–17, 01 Jan. 2015 2015, doi: 10.1123/JAPA.2013-0103.

[78] T. Radtke, M. Rodriguez, J. Braun, and H. Dressel, “Criterion validity of the ActiGraph and activPAL in classifying posture and motion in office-based workers: A cross-sectional laboratory study,” (in eng), Plos One, vol. 16, no. 6, p. e0252659, 2021, doi: 10.1371/journal.pone.0252659.

[79] C. L. Edwardson et al., “Considerations when using the activPAL monitor in field-based research with adult populations,” (in eng), J Sport Health Sci, vol. 6, no. 2, pp. 162–178, Jun 2017, doi: 10.1016/j.jshs.2016.02.002.

[80] E. S. Izmailova et al., “Implementing sensor-based digital health technologies in clinical trials: Key considerations from the eCOA Consortium,” (in eng), Clin Transl Sci, vol. 17, no. 11, p. e70054, Nov 2024, doi: 10.1111/cts.70054.

[81] G. Mitsi et al., “Implementing Digital Technologies in Clinical Trials: Lessons Learned,” (in eng), Innov Clin Neurosci, vol. 19, no. 4-6, pp. 65–69, Apr-Jun 2022.

[82] P. C. Fino et al., “Abnormal Turning and Its Association with Self-Reported Symptoms in Chronic Mild Traumatic Brain Injury,” (in English), J Neurotraum, vol. 35, no. 10, pp. 1167–1177, May 2018, doi: 10.1089/neu.2017.5231.

[83] G. E. Grossman, R. J. Leigh, L. A. Abel, D. J. Lanska, and S. E. Thurston, “Frequency and velocity of rotational head perturbations during locomotion,” Experimental Brain Research, vol. 70, no. 3, pp. 470–476, 1988/05/01 1988, doi: 10.1007/BF00247595.

[84] A. R. Weston, B. J. Loyd, C. Taylor, C. Hoppes, and L. E. Dibble, “Head and Trunk Kinematics during Activities of Daily Living with and without Mechanical Restriction of Cervical Motion,” (in eng), Sensors (Basel), vol. 22, no. 8, Apr 16 2022, doi: 10.3390/s22083071.

[85] A. Ashburn, C. Kampshoff, M. Burnett, E. Stack, R. Pickering, and G. Verheyden, “Sequence and onset of whole-body coordination when turning in response to a visual trigger: comparing people with Parkinson’s disease and healthy adults,” Gait Posture, vol. 39, no. 1, pp. 278–283, 2014.

[86] P. Crenna et al., “The association between impaired turning and normal straight walking in Parkinson’s disease,” Gait Posture, vol. 26, no. 2, pp. 172–178, 2007.

[87] F. Huxham, R. Baker, M. E. Morris, and R. Iansek, “Head and trunk rotation during walking turns in Parkinson’s disease,” Movement disorders: official journal of the Movement Disorder Society, vol. 23, no. 10, pp. 1391–1397, 2008.

[88] J. E. Visser et al., “Quantification of trunk rotations during turning and walking in Parkinson’s disease,” Clinical Neurophysiology, vol. 118, no. 7, pp. 1602–1606, 2007.

[89] D. Q. McDonald et al., “Head Movement Patterns during Face-to-Face Conversations Vary with Age,” presented at the Companion Publication of the 2022 International Conference on Multimodal Interaction, Bengaluru, India, 2022. [Online]. Available: 10.1145/3536220.3563366.

[90] K. B. Martin et al., “Objective measurement of head movement differences in children with and without autism spectrum disorder,” Molecular autism, vol. 9, pp. 1–10, 2018.

[91] E. Z. McClave, “Linguistic functions of head movements in the context of speech,” Journal of pragmatics, vol. 32, no. 7, pp. 855–878, 2000.

[92] Z. Zhao et al., “Atypical head movement during face‐to‐face interaction in children with autism spectrum disorder,” Autism Research, vol. 14, no. 6, pp. 1197–1208, 2021.

[93] Z. Zhao et al., “Identifying autism with head movement features by implementing machine learning algorithms,” Journal of Autism and Developmental Disorders, pp. 1–12, 2022.

[94] C. H. Meyer, A. G. Lasker, and D. A. Robinson, “The upper limit of human smooth pursuit velocity,” Vision research, vol. 25, no. 4, pp. 561–563, 1985.

[95] C. Tudor-Locke et al., “How many steps/day are enough? For older adults and special populations,” (in eng), Int J Behav Nutr Phys Act, vol. 8, p. 80, Jul 28 2011, doi: 10.1186/1479-5868-8-80.

[96] G. Baker et al., “The effect of a pedometer-based community walking intervention “Walking for Wellbeing in the West” on physical activity levels and health outcomes: a 12-week randomized controlled trial,” (in eng), Int J Behav Nutr Phys Act, vol. 5, p. 44, Sep 5 2008, doi: 10.1186/1479-5868-5-44.

[97] A. Weiss et al., “Does the Evaluation of Gait Quality During Daily Life Provide Insight Into Fall Risk? A Novel Approach Using 3-Day Accelerometer Recordings,” Neurorehabilitation and Neural Repair, vol. 27, no. 8, pp. 742–752, 2013, doi: 10.1177/1545968313491004.

[98] J. Spildooren, C. Vinken, L. Van Baekel, and A. Nieuwboer, “Turning problems and freezing of gait in Parkinson’s disease: a systematic review and meta-analysis,” (in English), Disabil Rehabil, vol. 41, no. 25, pp. 2994–3004, 2019, doi: 10.1080/09638288.2018.1483429.

